# The role of indirect effects in coevolution as mutualism transitions into antagonism

**DOI:** 10.1101/2021.11.22.469544

**Authors:** Fernando Pedraza, Hanlun Liu, Klementyna A. Gawecka, Jordi Bascompte

## Abstract

Species interactions have evolved from antagonistic to mutualistic and back several times throughout life’s history. Yet, it is unclear how changes in the type of interaction between species alter the coevolutionary dynamics of entire communities. This is a pressing matter, as transitions from mutualisms to antagonisms may be becoming more common with human-induced global change. Here, we combine network and evolutionary theory to simulate how shifts in interaction types alter the coevolution of empirical communities. We show that as mutualistic networks shift to antagonistic, selection imposed by direct partners begins to outweigh that imposed by indirect partners. This weakening of indirect effects is associated with communities losing their tight integration of traits and increasing their rate of adaptation. The above changes are more pronounced when specialist consumers are the first species to switch to antagonism. A shift in the outcome of species’ interactions may therefore reverberate across communities and alter the direction and speed of coevolution.

## Introduction

Mutualisms have arisen from antagonistic interactions multiple times throughout the history of life [1, 2, 3, 4]. Although no single mechanism explains these transitions, they are, to an extent, a result of changes in the costs and benefits of the interactions [5, 6, 7]. These cost-benefit relations are, in turn, shaped by biotic and abiotic factors [7]. Just as some conditions may ease the transition from antagonism to mutualism, others can reverse the transition and break mutualisms down [8, 9]. For example, cheating can arise in arbuscular mycorrhizal interactions [10], parasitism can overtake specialized and strict mutualisms like fig-wasp interactions [11], while fruit–frugivore interactions may switch from seed dispersal to seed predation [12]. There is also increasing evidence of shifts from mutualism to antagonism due to human-induced global change [9]. For example, ants have been found to switch to an antagonistic relationship with *Acacia* trees in the absence of large herbivores [13], while invasive pollinators can damage flowers impacting plant reproduction [14, 15]. The fact that the outcome of species interactions are context dependent [5, 6, 7], suggests that mutualisms and antagonisms are part of a continuum [16, 17].

Despite the well-documented examples of transitions in interaction types, their evolutionary consequences are not yet fully understood in the context of ecological communities. Since the coevolutionary dynamics of a community depend on the interaction type [18], a switch from mutualism to antagonism will lead to altered reciprocal evolutionary change of interacting species. Moreover, since coevolution operates through direct and indirect interactions at the community level [19, 20, 21], the effect of an interaction switching from mutualistic to antagonistic could in fact ripple across the community. Yet, it is unclear to what extent the coevolutionary dynamics typical of mutualistic systems, e.g. strong contribution of indirect effects to coevolution, high trait matching between partners, and a slow rate of adaptive change [21], may be derailed with a shift to antagonism.

Here, we combine network and evolutionary theory to simulate the evolutionary consequences of a shift from mutualistic to antagonistic interactions in communities. Specifically, we focus on measuring how the role of indirect effects in shaping coevolution changes in networks with increasing fractions of antagonistic interactions. We investigate how higher levels of antagonism alter the relationships between the contribution of indirect effects to coevolution, network structure, and trait distributions. Furthermore, we explore these trends under two scenarios: one where the shift to antagonism begins with specialist species, and the other with generalists.

## Methods

Our workflow consisted of five stages. First, we modified a set of empirical mutualistic networks to include different ratios of antagonistic-to-mutualistic interactions (see Figure 1). Second, we simulated species coevolution on the networks. Third, for each network, we measured its structural properties, the role of indirect effects on species coevolution, and the degree of trait matching arising after coevolution. Fourth, for each species, we measured its number of interactions (i.e. degree), contribution to indirect effects and degree of trait matching after coevolution. Fifth, we analysed how indirect effects and trait matching changed along the different ratios of antagonistic interactions, and across network structures.

**Figure 1:**
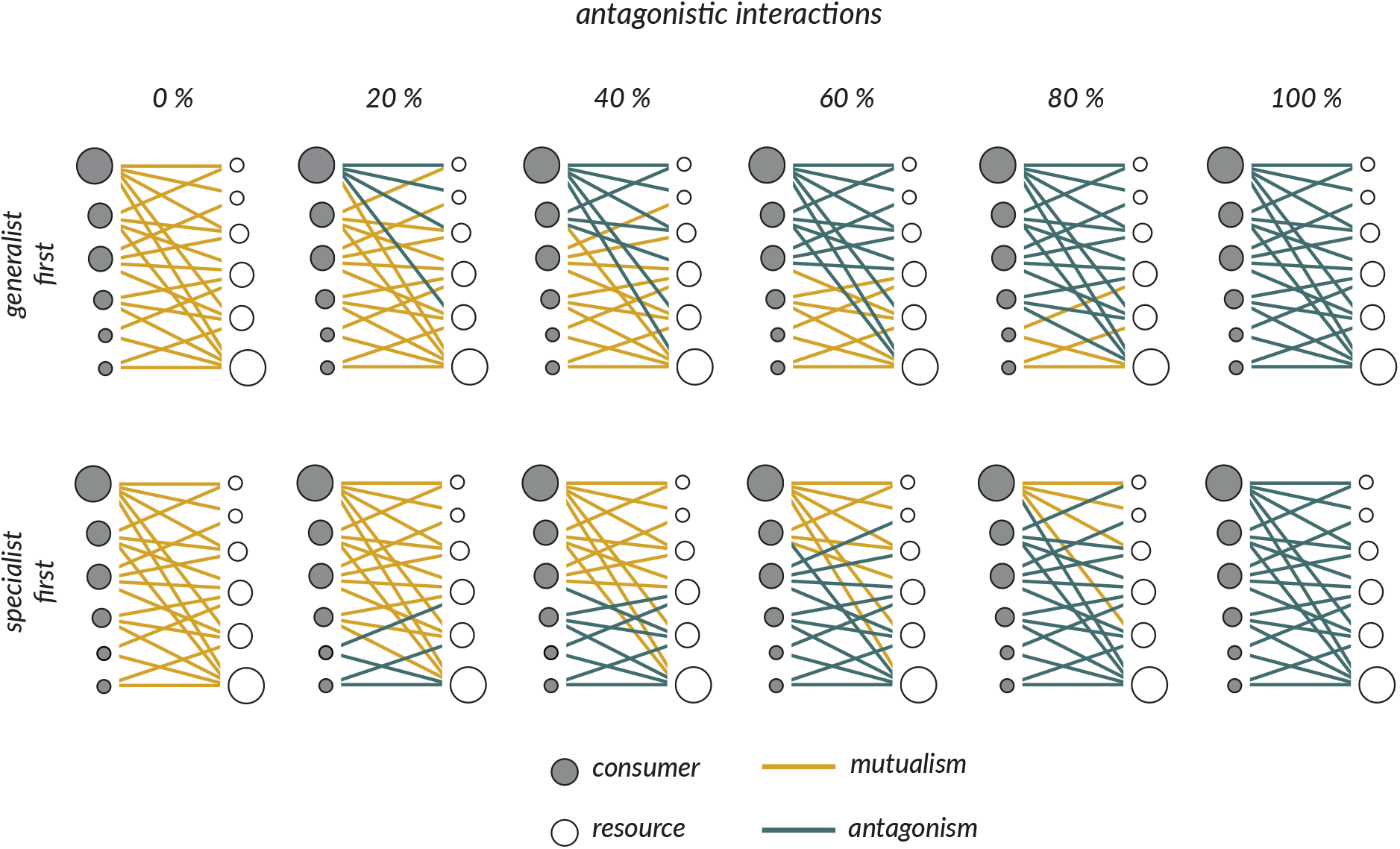
The shift from mutualistic to antagonistic networks. The figure depicts how we transform a mutualistic network (leftmost network) to a fully antagonistic one (rightmost network). At each fraction of antagonistic interactions we convert the corresponding number of links from mutualistic to antagonistic following two strategies. The ‘generalist first’ strategy (top panel) converts the links of the consumer species with the largest number of interactions first. The ‘specialist first’ strategy (bottom panel) converts the links of the consumer species with the least number of interactions first. At a given fraction of antagonistic interactions, both strategies result in a network with the same number of antagonistic interactions, but differ in how they are distributed. The networks resulting from each strategy are the same when there are 0 or 100% antagonistic interactions. The transformations do not alter network structure, they only change the interaction types.

### Dataset and conversion of interaction types

We used a set of mutualistic networks (seed dispersal networks, n = 34) found in the Web of Life repository (http://www.web-of-life.es/, [22]). In these networks, the species are linked by mutualistic interactions. To obtain networks with a mix of antagonistic and mutualistic interactions, we converted a proportion of links from mutualistic to antagonistic (for example, to obtain a network with 20% antagonistic interactions, we chose 20% of the links of the network and reassigned them as antagonistic interactions). These transformations did not alter network structure. They merely changed the type of link connecting species, with implications on how coevolution unfolded (see Section ‘Coevolution model and simulations’).

We transformed each mutualistic network to obtain an equivalent network with 0, 20, 40, 60, 80, and 100% of antagonistic interactions. To decide which interactions to transform first, we followed two different methods. Both strategies relied on measuring the number of interactions each consumer species had in the network. The ‘generalist first’ strategy converted the interactions of the consumer species with the highest degree in the network first (Figure 1, top panel). The ‘specialist first’ strategy prioritized the consumer species with the lowest degree in the network (Figure 1, bottom panel). Both strategies assume that any switch from mutualism to antagonism is driven by consumers. Note that it is possible for both strategies to happen in nature. On one hand, generalist species are sometimes opportunists with less specialized traits or are unreliable at rewarding their mutualistic partners [23, 24], which may make them more likely to transition to antagonism [25, 26]. On the other hand, the emergence of antagonistic cheaters is limited by disfavour or sanction from their partners, and therefore cheating may tend to occur in specialists with a few susceptible partners [27, 28, 10]. For completeness, we also converted interactions at random. However, as we found that the other two strategies represent the two extreme scenarios, and as the results from this ‘random’ strategy followed the same trends, they are not shown here.

To rule out the possibility that the trends we observed were driven by the structure of the mutualistic networks selected as opposed to the conversion of interactions to antagonistic, we also transformed a set of antagonistic networks (n = 55, 51 host-parasites and 4 plant-herbivore networks) from the Web of Life repository. In this case, we converted the originally antagonistic interactions into mutualistic ones and performed the same workflow as outlined above.

### Coevolution model and simulations

To simulate coevolution, we used a discrete model that uses a selection gradient approach to link trait evolution with the fitness consequences of interactions and the environment [21]. In essence, the model assigns an initial trait value to each individual of each species in the network and updates it over time as a result of selection imposed by partners and the environment. The evolution in discrete time of the mean trait value of the population of species *i* (*Z_i_*) can be expressed as:

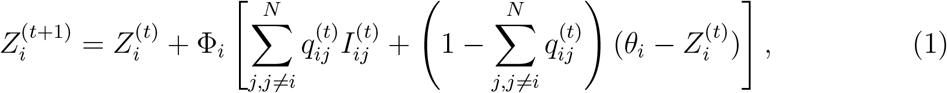

where Φ_*i*_ is proportional to the selection gradient and additive genetic variance, *N* is the number of species in the network, *q_ij_* is the evolutionary effect of species *j* on species *i*, where 0 ≤ *q_ij_* ≤ 1 (Eq. 2), *I_ij_* is the phenotype selected by the interaction of species *j* with *i*, and *θ_i_* is the environmental optimum for species *i*.

The evolutionary effects of species *j* on species *i* are defined by:

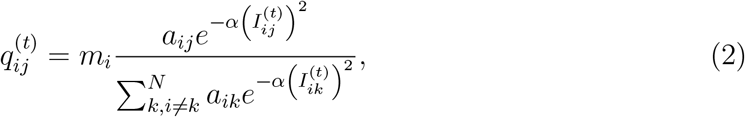

where *m_i_* is the level of coevolutionary selection 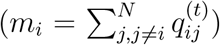, *α* is a constant that determines the sensitivity of the evolutionary effect to *I_ij_*, and *a_ij_* is an element of an adjacency matrix *A* of a given network. *a_ij_* = 1 if species *i* interacts with species *j*, and *a_ij_* = 0 if they do not. The coevolutionary selection parameter (0 ≤ *m_i_* ≤ 1) determines the relative importance of coevolution in driving trait evolution.

We assume that mutualistic and antagonistic interactions differ in how they affect trait evolution. That is, the phenotype selected by the interaction of species *j* with *i* (*I_ij_* in Eq. 1 and Eq. 2) will depend on the type of interaction. Following Guimarães et al. [21], we assume that mutualistic interactions favour trait similarity between partners, regardless of the guild the species belongs to. Hence, the effect of species *j* on species *i* at time *t* can be expressed as:

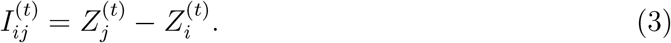

Following Andreazzi et al. [29, 30] we assume that antagonistic interactions favour trait similarity on consumers and mismatch on resource species. From the consumer perspective, the effect of resource *j* on consumer *i* at time *t* is defined by Eq. 3. Trait mismatch will result in resource species’ traits either increasing (if *Z_resource_* > *Z_consumer_*) or decreasing (if *Z_resource_* < *Z_consumer_*). As in Andreazzi et al. [29, 30], we define a critical mismatch value *ε*. If |*Z_consumer_* – *Z_resource_*|> *ε*, then the consumer species has a negligible effect on the resource (*q_i_j* = 0). From the resource perspective, the effect of consumer *j* on resource *i* at time t can be expressed as:

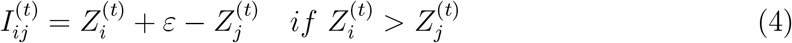

or

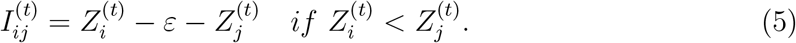

We used the coevolution model (Eq. 1) to simulate coevolution on all networks at each particular fraction of antagonistic interactions (i.e. 0, 20, 40, 60, 80, and 100%). We sampled the values for *θ, Φ, m* and the initial trait values for all species 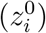 from statistical distributions at the start of each simulation (see Table S1). We ran simulations until equilibrium, defined as 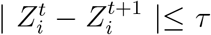, with *τ* = 1e-5 and *α* = 0.2 [21]. We ran 100 replicates per network for a total of 40,800 simulations (34 networks × 6 fractions × 2 strategies). Each replicate was assigned different parameter values which were then kept the same across all fractions of antagonistic interactions.

### Data analysis

#### Network structure

We measured four structural properties of the networks we used in this study. These were, network size (s), connectance (c), modularity (M), and nestedness (N). To identify modules and quantify the modularity of a network, we used the *Q* metric [31, 32]. *Q* measures the difference between the observed fraction of interactions between species in the same module and the expected fraction of interactions connecting species in the same module if interactions were established at random. We used the MODULAR software [32] with a simulated annealing algorithm [33] to find the network partition where *Q* is maximized, and recorded the *Q*-value of the network partition. Nestedness is a measure of the extent to which more specialist species only interact with subsets of those species interacting with more generalist species [34]. We quantified network nestedness using a metric proposed by Fortuna et al. [35], which is equivalent to the NODF metric [36]. This allowed us to measure nestedness as the average overlap between interactions of consumers (resources), without penalizing the contribution to nestedness of consumers (resources) able to interact with the same number of resources (consumers). Lastly, we measured the number of interactions (i.e. degree) of each species in the networks.

We summarized all four network descriptors into a single metric by means of a principal component analysis (PCA). We described the structure of each network by extracting its position along the first component (PC1) of the PCA. PC1 explained 75% of the variance in structural descriptors and was negatively correlated with network size (−0.65) and modularity (−0.88), and positively correlated with connectance (0.94) and nestedness (0.94).

#### Trait matching

We calculated the degree of trait matching between all pairs of species in the network when simulations reached equilibrium. We defined trait matching (*M*) between a pair of species *i, j* at a given time *t* as:

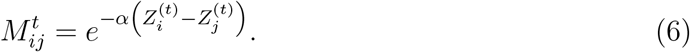

We used Eq. 6 to measure the mean trait matching value for each network and for each species in the network. For each network and species, we calculated the average trait matching across replicates. Thus, trait matching is a summary measure of all pairwise trait differences that allows us to make comparisons across species and networks. We relate the level of trait matching at the network (or species) level with network structure (or species’ degree).

#### Indirect effects

We measured the contribution of indirect effects to trait evolution using the coevolutionary matrix (T-matrix) that arose when simulations reached equilibrium (see [21] for a detailed description). It is defined as:

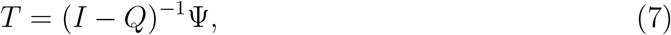

where *i* is the identity matrix, *Q* is an *N* x *N* matrix containing the direct coevolutionary effects of interactions (Eq. 2), and Ψ is an *N* x *N* diagonal matrix with Ψ_*ii*_ = 1 – *m_i_* for species *i* (note that as in the coevolution model, m represents the level of coevolutionary selection). In short, the T-matrix contains the relative contribution of each interacting and non-interacting species in the network to the selection gradient shaping trait evolution of a given species in the network [21]. We used the elements of the T-matrix (*t_ij_*) and adjacency matrix (*a_ij_*) of a network to extract the contribution of non-interacting species on the trait evolution of each species in the network. We then defined the relative contribution of indirect effects to trait evolution in a network as:

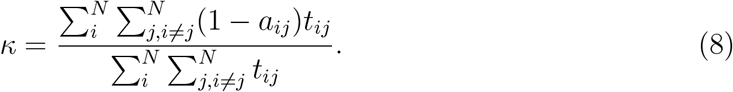

In addition, we measured the contribution of each species to the proliferation of indirect effects in the network. For species *i*, this is defined as:

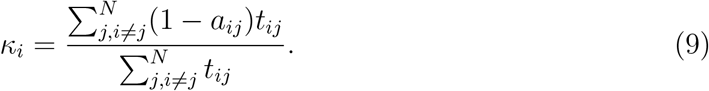

Note that, the contribution of indirect effects is related to both the network structure (through the adjacency matrix), and the degree of trait matching (through matrix Q and Eq. 2). For each network and species, we calculated the average contribution of indirect effects to trait evolution across replicates. We relate the contribution of indirect effects to coevolution at the network (or species) level with network structure (or species’ degree).

#### Statistical analyses

We used simple linear regressions to determine whether the relationship between network structure (network coordinates on PC1) and indirect effects or trait matching changed with the fraction of antagonistic interactions in the network. We fit a single linear model for each fraction of antagonistic interactions with indirect effects (or trait matching) as the response variable and network structure as the explanatory variable. For each model, we extracted the estimated slope coefficients and their respective confidence intervals (95%). We performed all simulations, analyses, and visualizations in R version 4.0.2 [37].

## Results

We found that higher fractions of antagonistic interactions reduced the contribution of indirect effects to trait evolution (Figure 2A) and increased the rate of adaptive change (i.e. the average amount of trait change per time step) in the communities (Figure 2B). The changes to coevolutionary dynamics were more drastic when specialist consumers were the first to switch to antagonism (Figure 2A-D). We observed the same trends regardless of whether we converted interactions to antagonistic in mutualistic networks (Figure 2) or converted interactions to mutualistic in antagonistic networks (Figure S1).

**Figure 2:**
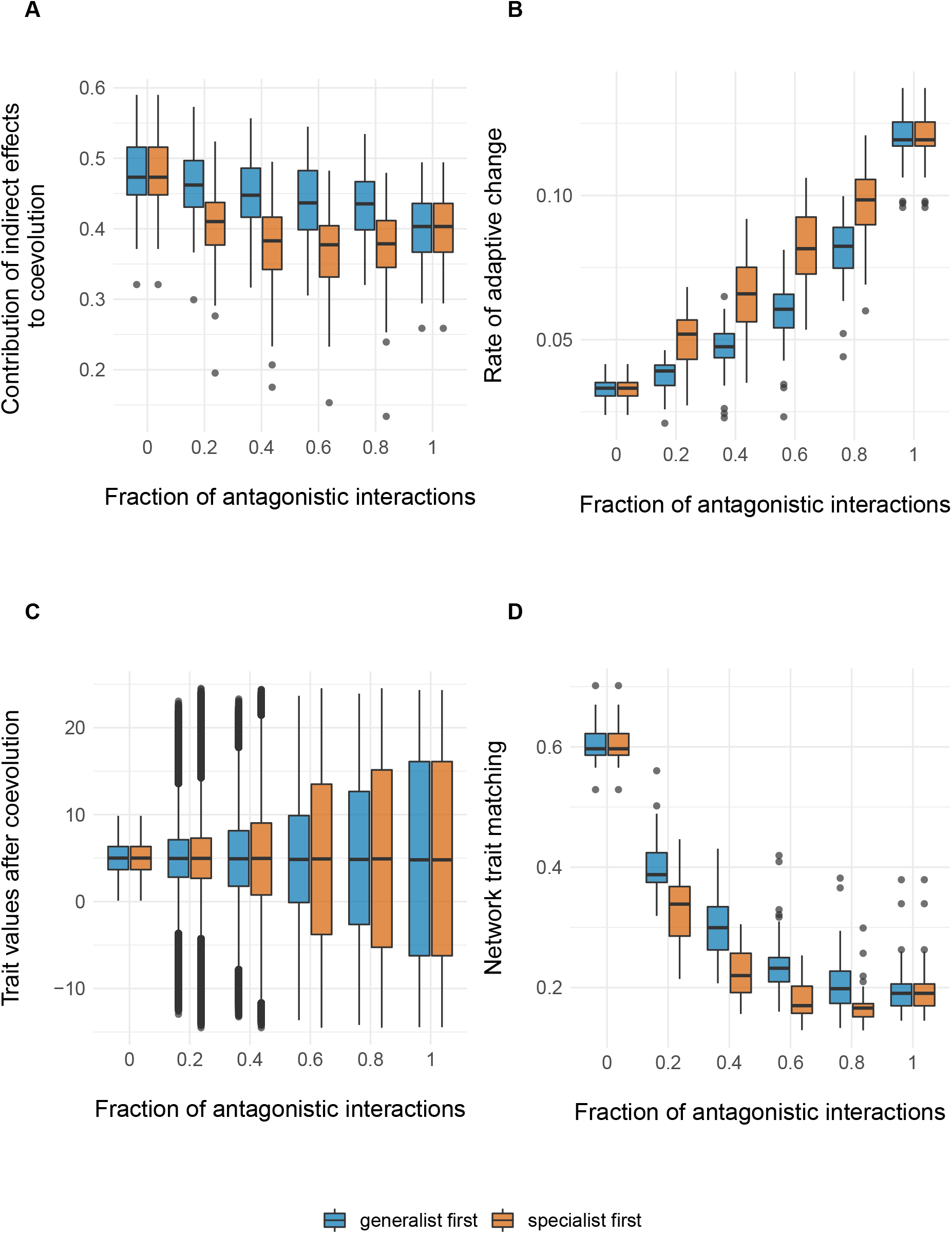
Evolutionary implications of a shift from mutualistic to antagonistic interactions. At a given fraction of antagonistic interactions, boxplots summarise the contribution of indirect effects to coevolution (A), the rate of adaptive change (B), the trait values (C), and the network trait matching (D) measured after simulating coevolution on a set of empirical networks (n = 34). Colours denote the strategy used to convert mutualistic to antagonistic interactions.

The reduction in the contribution of indirect effects to trait evolution could be explained by either (1) changes in how network structure shapes indirect effects or (2) alterations in the trait distributions of the communities. Next, we look at which one of these mechanisms explains our results.

The transition from mutualism to antagonism did not change how network structure affects indirect effects (or trait matching, Figures S2 and S3). At any given fraction of antagonistic interactions, smaller, more connected networks had lower contributions of indirect effects (and higher trait matching), than larger, more modular networks (Figure S2). We found the same associations between network structure and indirect effects after including networks where we converted antagonistic interactions to mutualistic (Figures S4 and S5). Thus, the diminishing role of indirect effects across the mutualism-to-antagonism continuum is not driven by a change in how network structure shapes indirect effects.

In contrast to the lack of effects of network structure, the shift from mutualism to antagonism did lead to changes in the trait distributions of the communities. As expected from our model, increasing the fraction of antagonistic interactions also increased the variance in trait values (Figure 2C) and, consequently, decreased trait matching in the network (Figure 2D). Surprisingly, these trends appear to be nonlinear, with the largest changes in trait matching occurring when the first 20% of interactions shift to antagonism. These observations held regardless of whether we converted interactions to antagonistic in mutualistic networks (Figure 2) or converted interactions to mutualistic in antagonistic networks (Figure S1).

The order in which a network transitioned from mutualism to antagonism affected the trait distributions of the entire community. We found that both consumers and resource species had greater variation in traits when specialist consumers switched to antagonism compared to when generalist consumers did so (Figure 3, differences between strategies for a given guild). Furthermore, at low fractions of antagonistic interactions, a consumer’s trait matching decreased with its degree under the ‘generalist first’ strategy, while it increased for the ‘specialist first’ strategy (Figure 4 0.2 - 0.6 consumer panel). For resource species, we observed the same general trends, but the strategies that generated them were the opposite to those described above (Figure 4 resource panel). In other words, if generalist consumers switch to antagonism first, then consumer trait matching decreases with degree, while resource trait matching increases with degree. Conversely, if specialist consumers become antagonistic first, then consumer trait matching increases with degree, while resource trait matching decreases with degree. Yet, note that the differences between strategies diminished as the fraction of antagonistic interactions increased. We observed similar trends when converting interactions to mutualistic in antagonistic networks (Figures S6 and S7).

**Figure 3:**
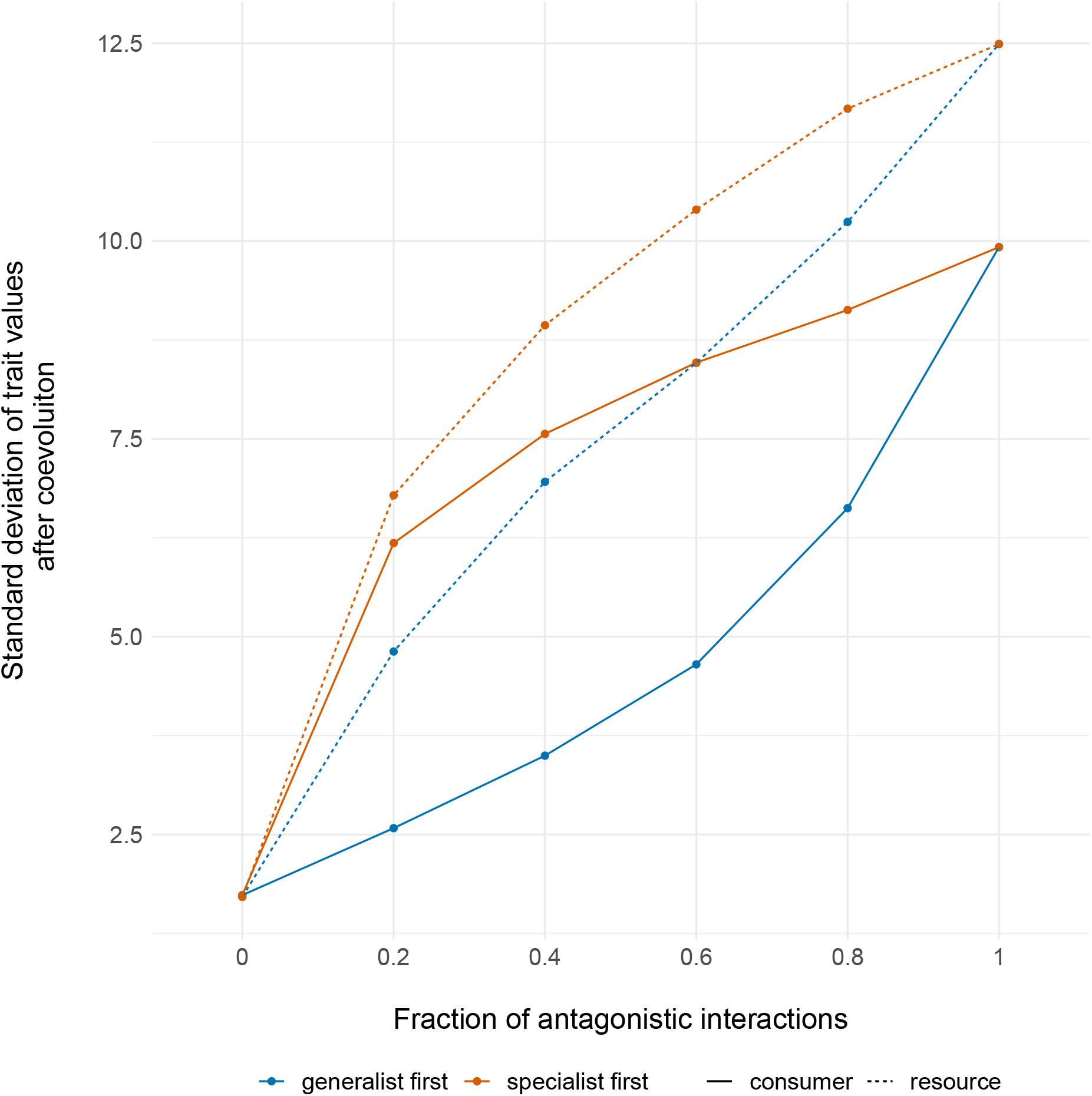
The spread of trait values in consumer and resource species as mutualistic interactions transition to antagonistic ones. Colours denote the strategy used to convert mutualisms to antagonistic interactions, while line types denote the guild (consumers or resources).

**Figure 4:**
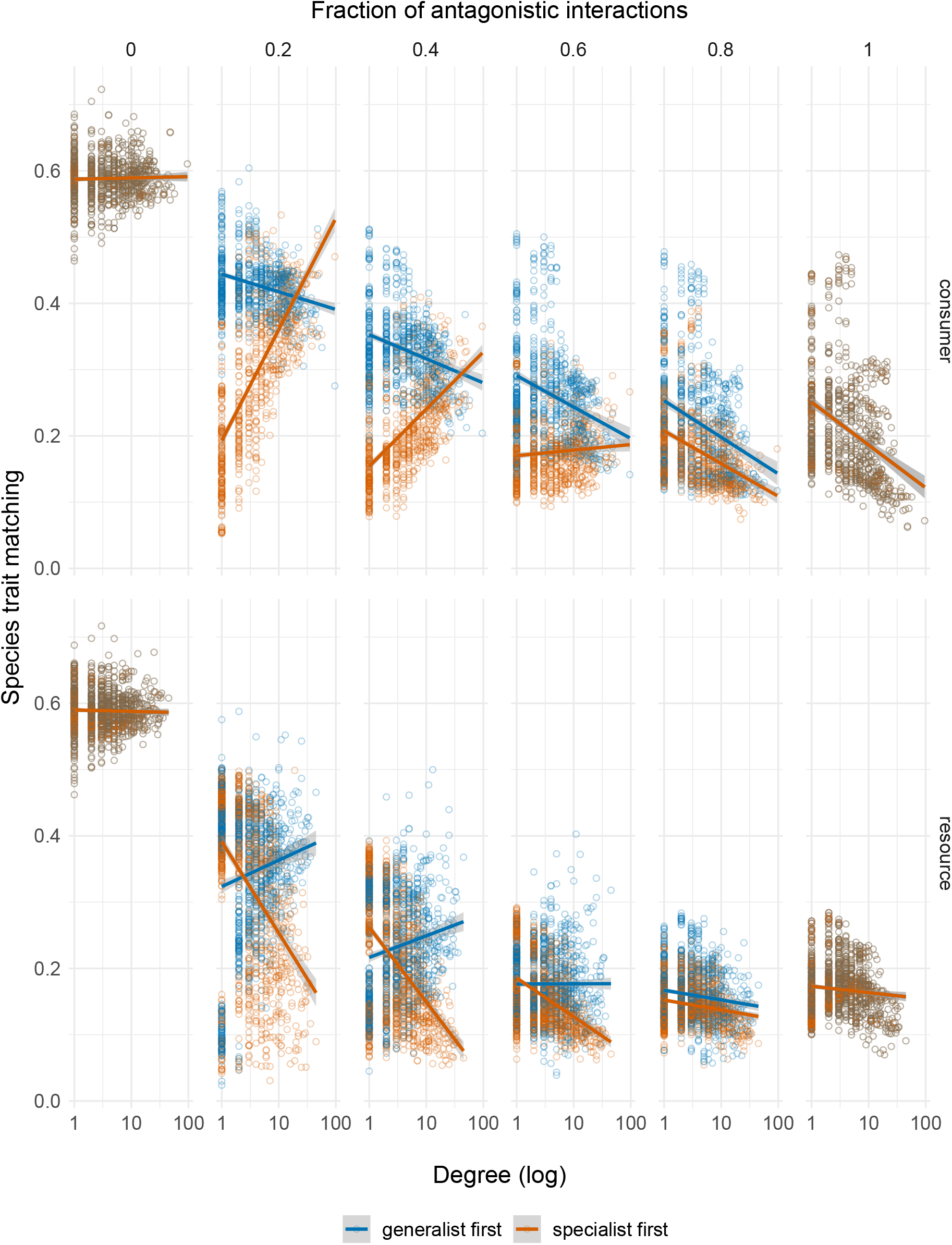
Effect of species degree on species trait matching, along the transition from mutualism to antagonism. Numbers above the panels denote the fraction of antagonistic interactions in the networks. Colours represent the strategy used to convert mutualisms to antagonistic interactions. Lines highlight the linear relation between variables.

## Discussion

Two main insights emerge from our work on the coevolutionary consequences of the shifts from mutualisms to antagonisms. First, the contribution of indirect effects to trait evolution is greater in mutualisms than antagonisms. Second, the influence of indirect effects on coevolution is lower if the shift to antagonism begins with specialist species. We argue that these observations are a result of changes in trait patterns arising from coevolution.

Why are indirect effects less influential in antagonism than mutualism? Since we used the same set of networks across the different fractions of antagonistic interactions, the changes in the contribution of indirect effects we observe are not associated with differences in network structure, but with the changes in trait distributions and matching. Three results suggest that indirect effects are positively associated with trait matching between species. First, under the ‘generalist first’ strategy, increasing the fraction of antagonistic interactions leads to an increase in trait spread, and a reduction of both trait matching and the contribution of indirect effects (Figure 2A, C and D). Second, for the ‘specialist first’ strategy, the contribution of indirect effects and trait matching jointly decrease until antagonistic interactions reach 80%, which is then followed by an increase in both variables with additional transitions to antagonism (Figure 2A and D). Third, when comparing species with the same degree, those that achieve higher trait matching have a higher contribution to indirect effects (Figures 4, S8 and S9). In summary, indirect effects contribute more to coevolution in mutualisms where species’ traits are more similar to each other.

Why are indirect effects less influential when the shift to antagonism begins with specialist species? Since our method of converting interactions from mutualistic to antagonistic does not alter species degree, differences in the contributions to indirect effects are driven by changes in trait values. The ‘specialist first’ strategy results in a larger trait spread and lower trait matching than the ‘generalist first’ strategy (Figure 2A, C and D). It follows that indirect effects are less influential in the ‘specialist first’ strategy. The lower trait matching resulting from the ‘specialist first’ strategy may be explained by the asymmetric selection pressures acting on consumers and resources, and the stronger influence that specialists have in driving trait patterns (Figure 4). From the consumer perspective, specialists evolve trait values closer to their few resource partners more easily than generalists, which have more partners. Thus, if specialist consumers switch to antagonism, they are likely to become more similar to their resource while becoming more dissimilar to the rest of the network. From the resource perspective, if generalists are targeted by specialist consumers, they evolve trait values that are more dissimilar not only to their consumers but to all other species. Taken together, these results highlight the importance of the role specialists play in shaping traits within networks and mediating species’ contribution to indirect effects.

It is important to keep in mind that the trait changes we observe are, to an extent, a result of our assumption that antagonisms lead to trait mismatch between partners, though they could be driving trait evolution in other ways [18, 38]. Additionally, we used the same network structures in the transition from mutualisms to antagonisms, despite ample evidence of differences in network structure between interaction types [39, 34, 40]. The potential rewiring of networks as interaction types change [9, 41] could impact how structure shapes indirect effects and trait matching. However, the fact that we observe the same trends when converting mutualistic networks to antagonistic, and antagonistic networks to mutualistic, may suggest that rewiring would not alter the patterns we described. Despite its limitations, our approach allowed us to explore hypotheses that are challenging to address through empirical observations.

While indirect effects drive coevolution in mutualisms [21], we show they have a weaker role in antagonisms. In other words, the transition from mutualism to antagonism leads to selection imposed by direct partners outweighing that imposed by indirect partners. This may hinder the emergence of community-wide trait convergence, while accelerating the ‘slow and steady’ pace of coevolution found in mutualisms [21]. We show that coevolutionary dynamics are modified most drastically when only a few interactions shift to antagonism. Furthermore, the effects of these conversions could be amplified if specialist consumers shift from mutualism to antagonism. Our results are particularly timely, as anthropogenic drivers such as climate change, habitat transformation or invasive species have already been shown to cause transitions from mutualistic to antagonistic interactions [8, 9, 42, 43, 44]. Thus, conditions that alter the outcome of species interactions may modify the direction and speed of coevolution of entire communities.

## Acknowledgements

We thank the members of the Bascompte Lab for their insightful comments.

## Data availability

All code is available on Github: https://github.com/fp3draza/indirect_effects_transition_mutualism_antagonism

## Author contributions

FP, HL, KAG, and JB designed research. FP and HL performed research. FP and KAG analysed data. FP wrote a first draft of the manuscript, and all authors contributed substantially to the final draft.

---

**Table S1:**
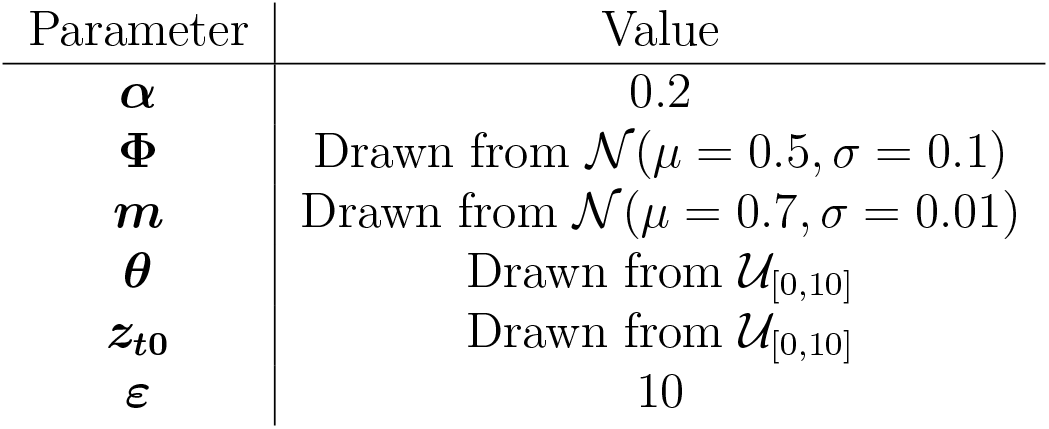
Values of the model parameters.

**Figure S1:**
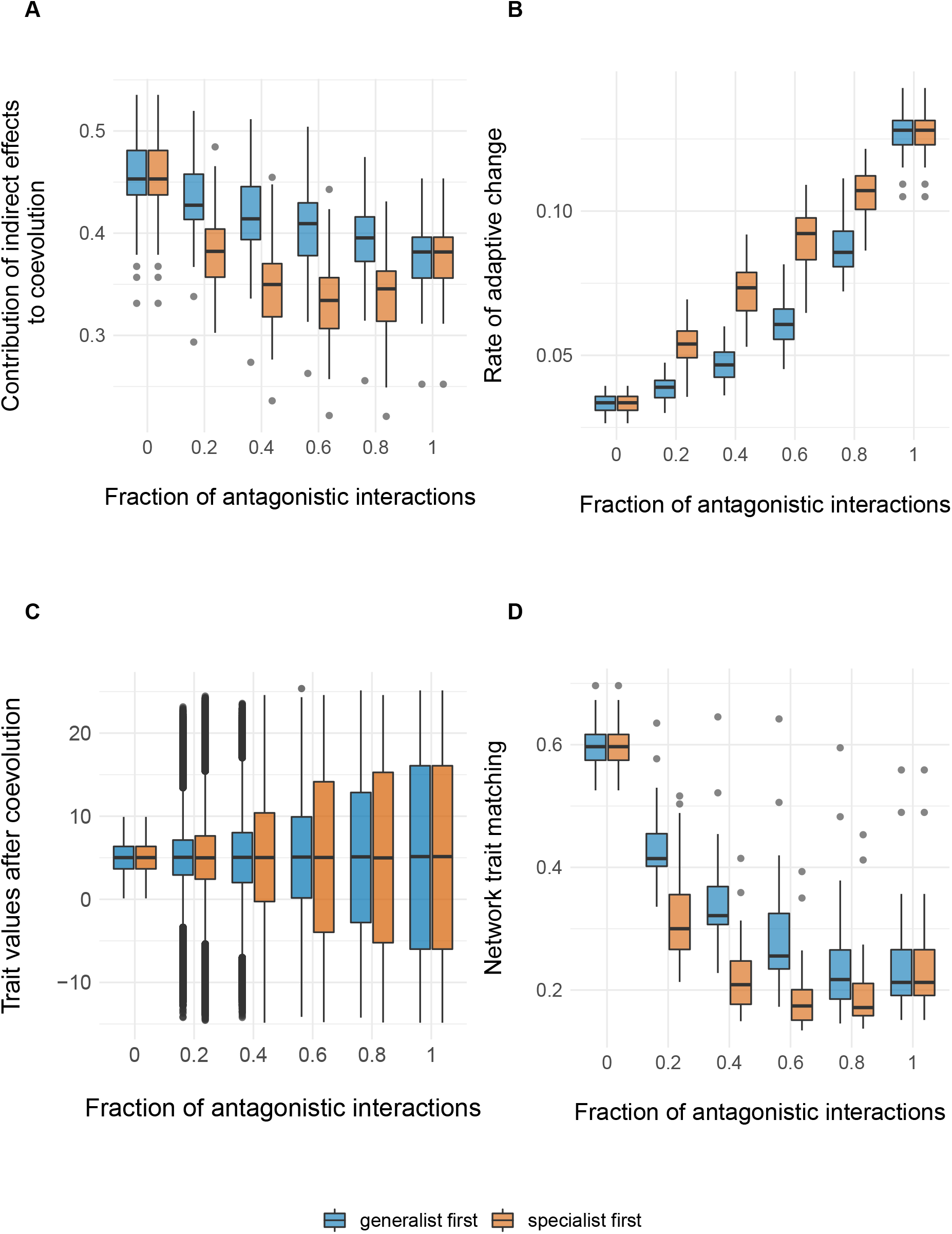
Evolutionary implications of a shift from antagonistic to mutualistic interactions. At a given fraction of antagonistic interactions, barplots summarise the contribution of indirect effects to coevolution (A), the rate of adaptive change (B), the trait values (C), and the network trait matching (D) measured after simulating coevolution on a set of empirical networks (n = 55). Colours denote the strategy used to convert antagonistic to mutualistic interactions.

**Figure S2:**
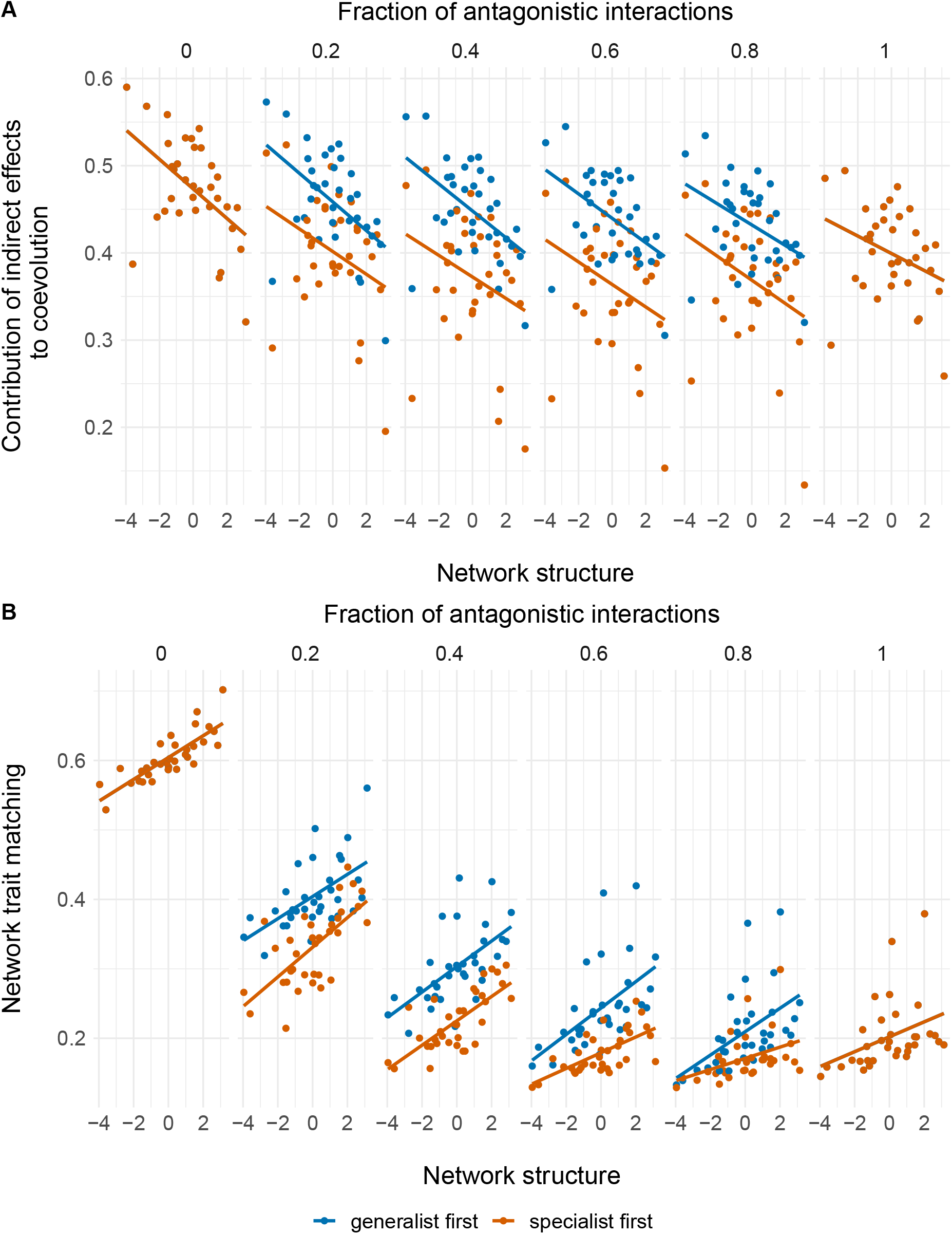
Effect of network structure on the contribution of indirect effects to coevolution (A), and trait matching (B). Points represent a network’s average indirect effects or trait matching after simulating coevolution. Positive values along the network structure axis denote a more connected, more nested network. Negative values denote a larger, more modular network. Colours denote the strategy used to convert mutualistic to antagonistic interactions.

**Figure S3:**
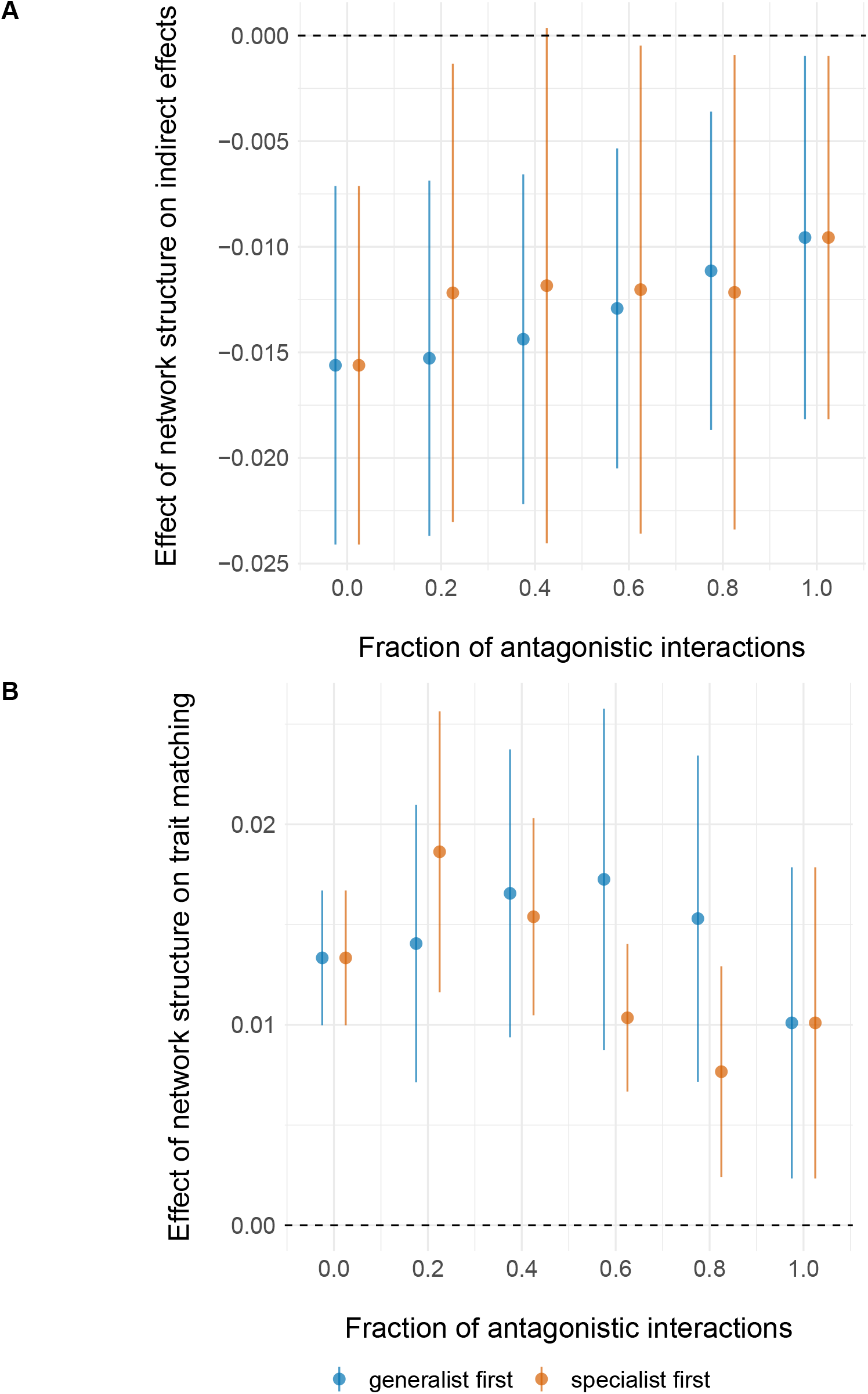
Effect of network structure across fractions of antagonistic interactions as estimated from the slope of a linear model. At a given fraction of antagonistic interactions, points represent the slope of the linear relationship between network structure and indirect effects (A) or trait matching (B). Lines show the confidence intervals (95%) of the estimate. Colours denote the strategy used to convert mutualisms to antagonistic interactions.

**Figure S4:**
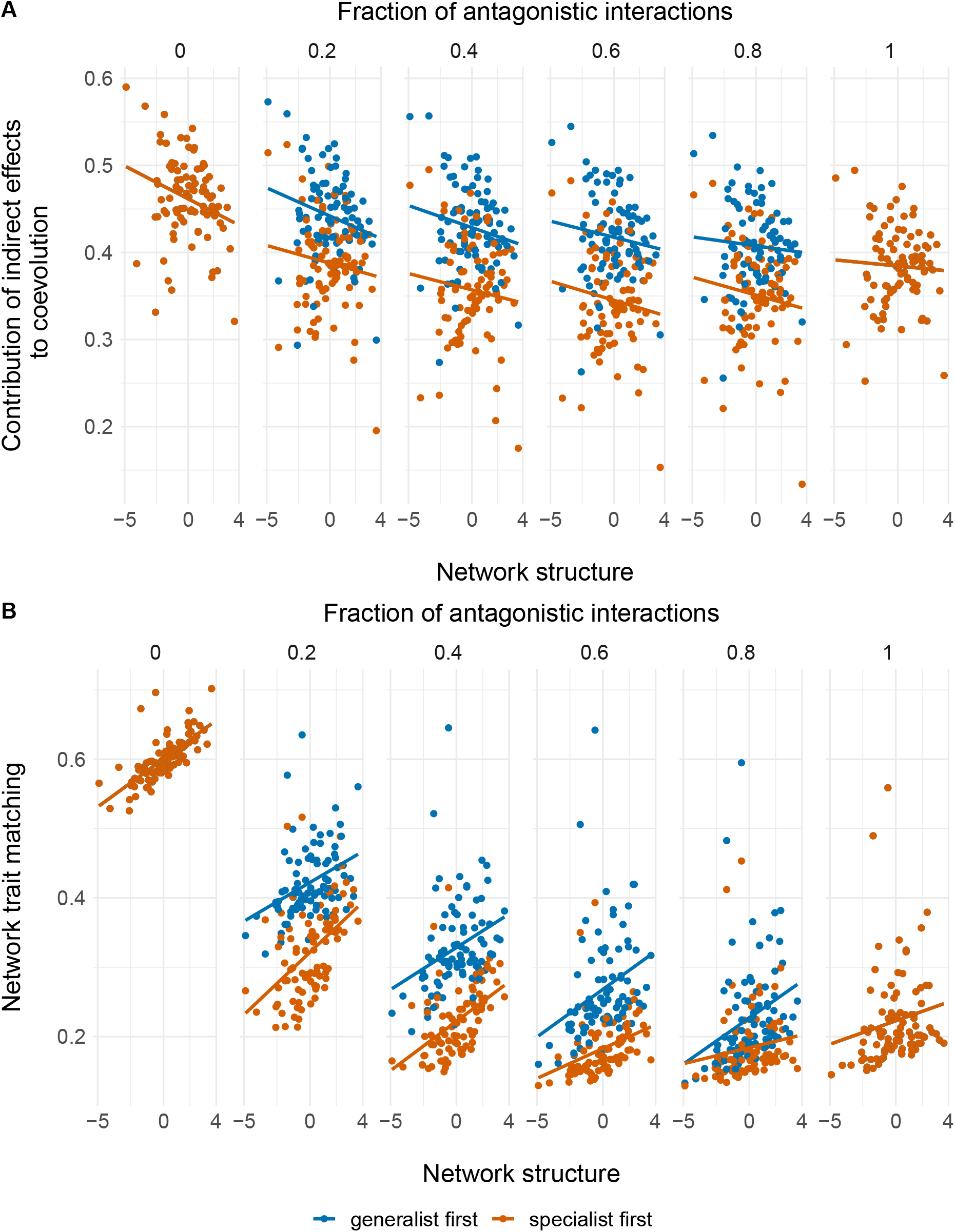
Effect of network structure on the contribution of indirect effects to coevolution (A), and trait matching (B). Points represent a network’s average indirect effects or trait matching after simulating coevolution. Positive values along the network structure axis denote a more connected, more nested network. Negative values denote a larger, more modular network. Colours denote the strategy used to convert antagonistic to mutualistic interactions.

**Figure S5:**
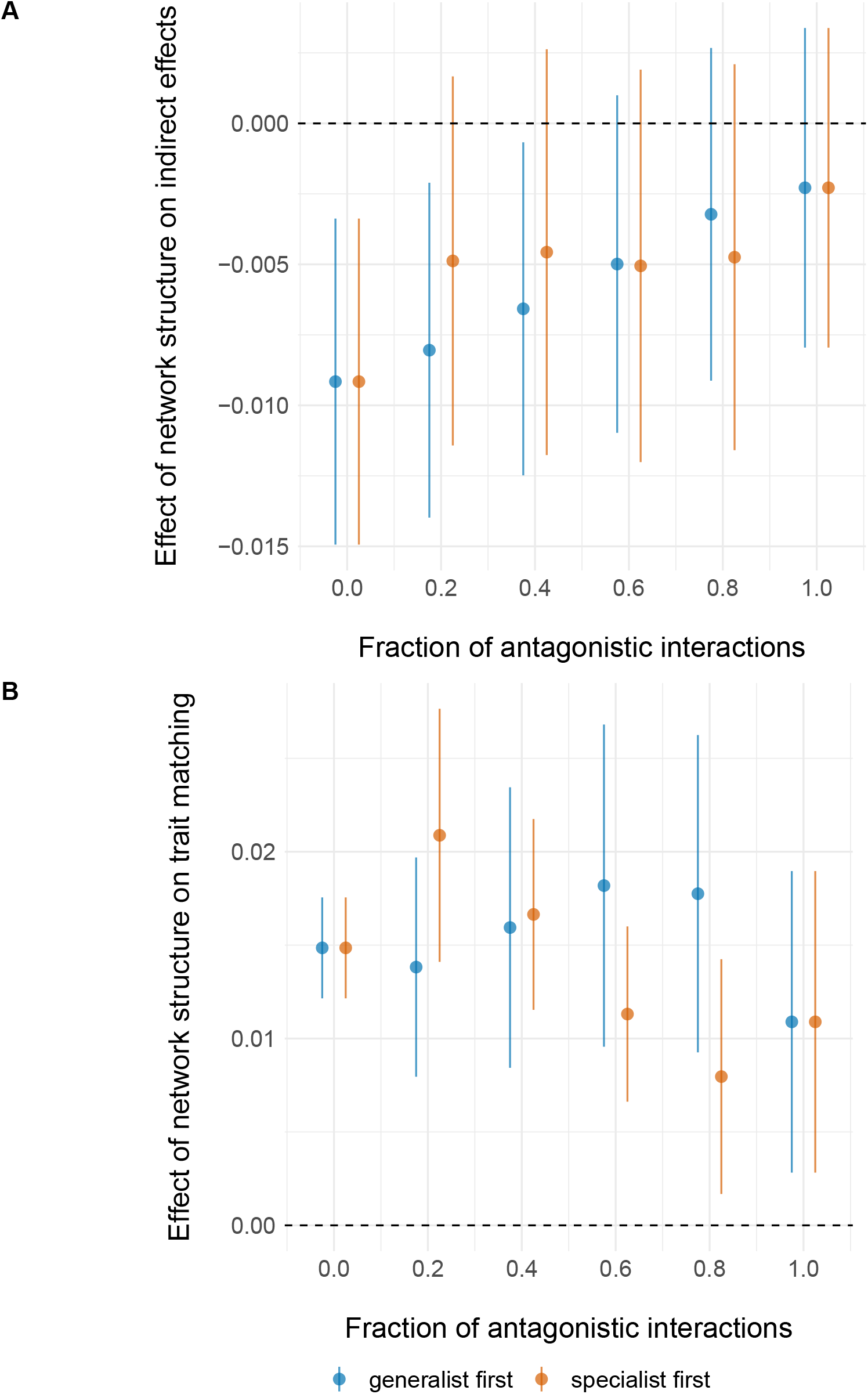
Effect of network structure across fractions of antagonistic interactions. At a given fraction of antagonistic interactions, points represent the slope of the linear relationship between network structure and indirect effects (A) or trait matching (B). Lines show the confidence intervals (95%) of the estimate. Colours denote the strategy used to convert antagonistic to mutualistic interactions.

**Figure S6:**
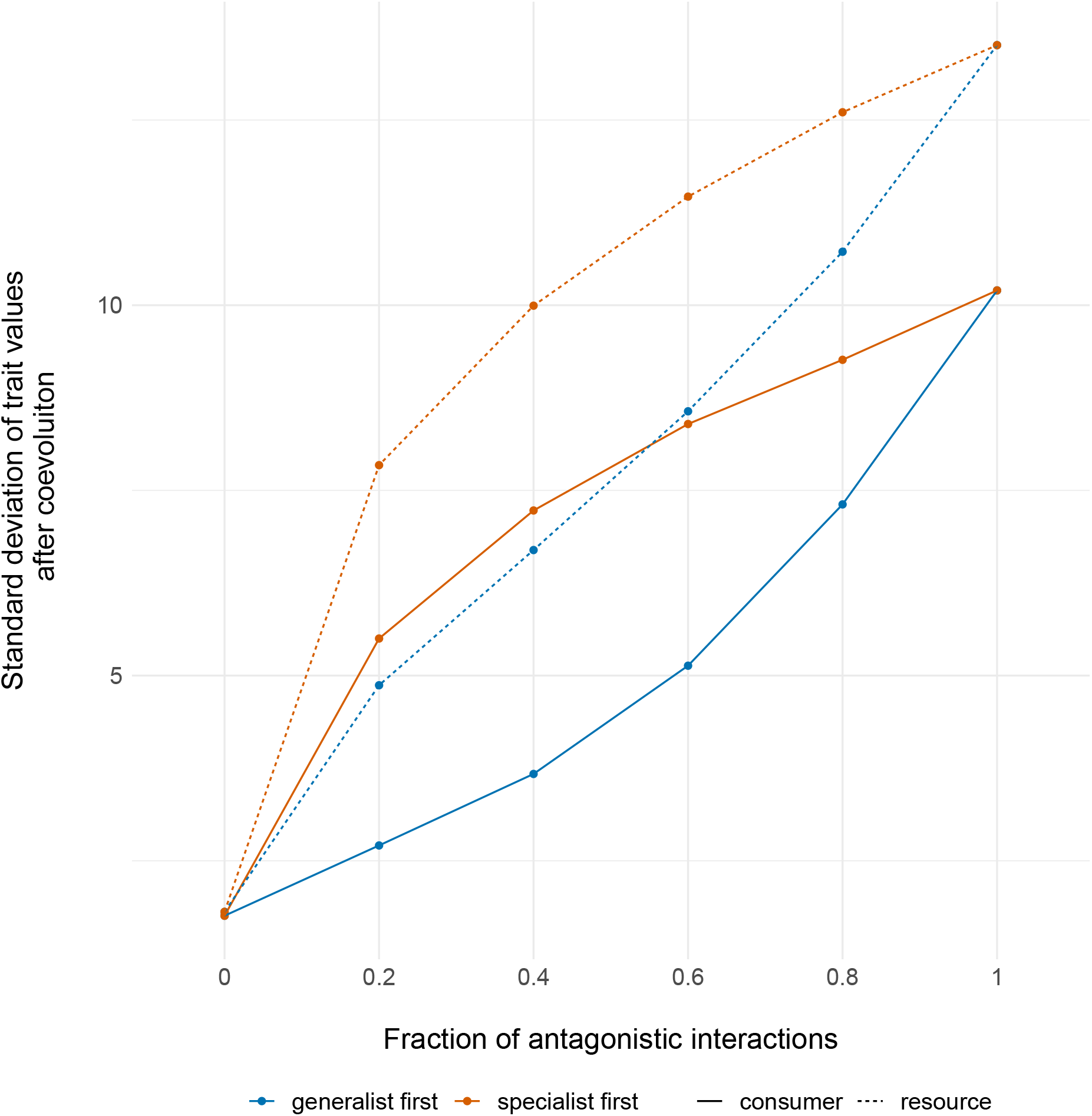
The spread of trait values in consumer and resource species as antagonistic interactions transition to mutualistic ones. Colours denote the strategy used to convert antagonistic to mutualistic interactions, while line types denote the guild (consumers or resources).

**Figure S7:**
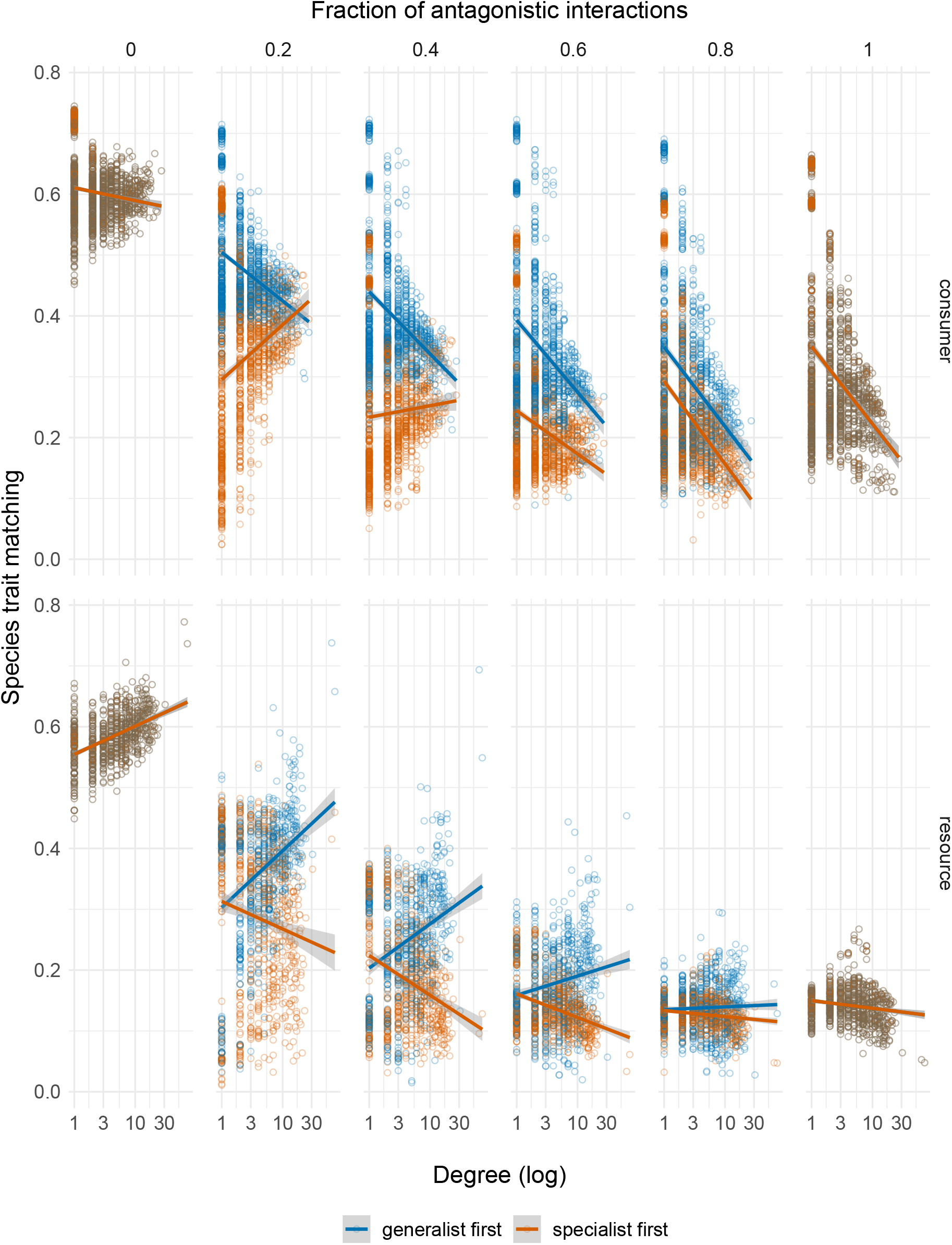
Effect of species degree on species trait matching, along the transition from antagonism to mutualism. Numbers above the panels denote the fraction of antagonistic interactions in the networks. Colours represent the strategy used to convert antagonistic to mutualistic interactions. Lines highlight the linear relation between variables.

**Figure S8:**
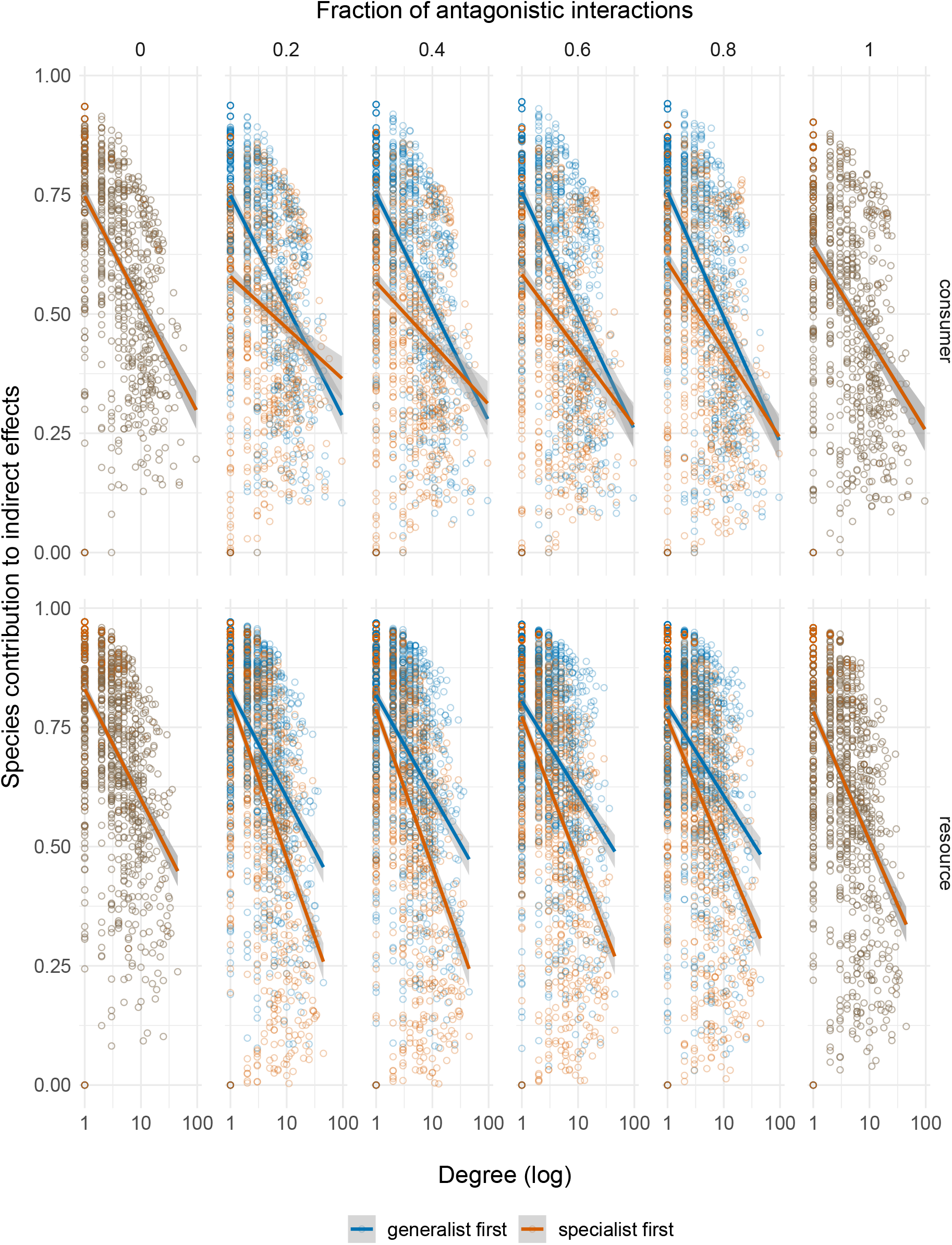
Effect of species degree on species contribution to indirect effects, along the transition from mutualism to antagonism. Numbers above the panels denote the fraction of antagonistic interactions in the networks. Colours represent the strategy used to convert mutualistic to antagonistic interactions. Lines highlight the linear relation between variables.

**Figure S9:**
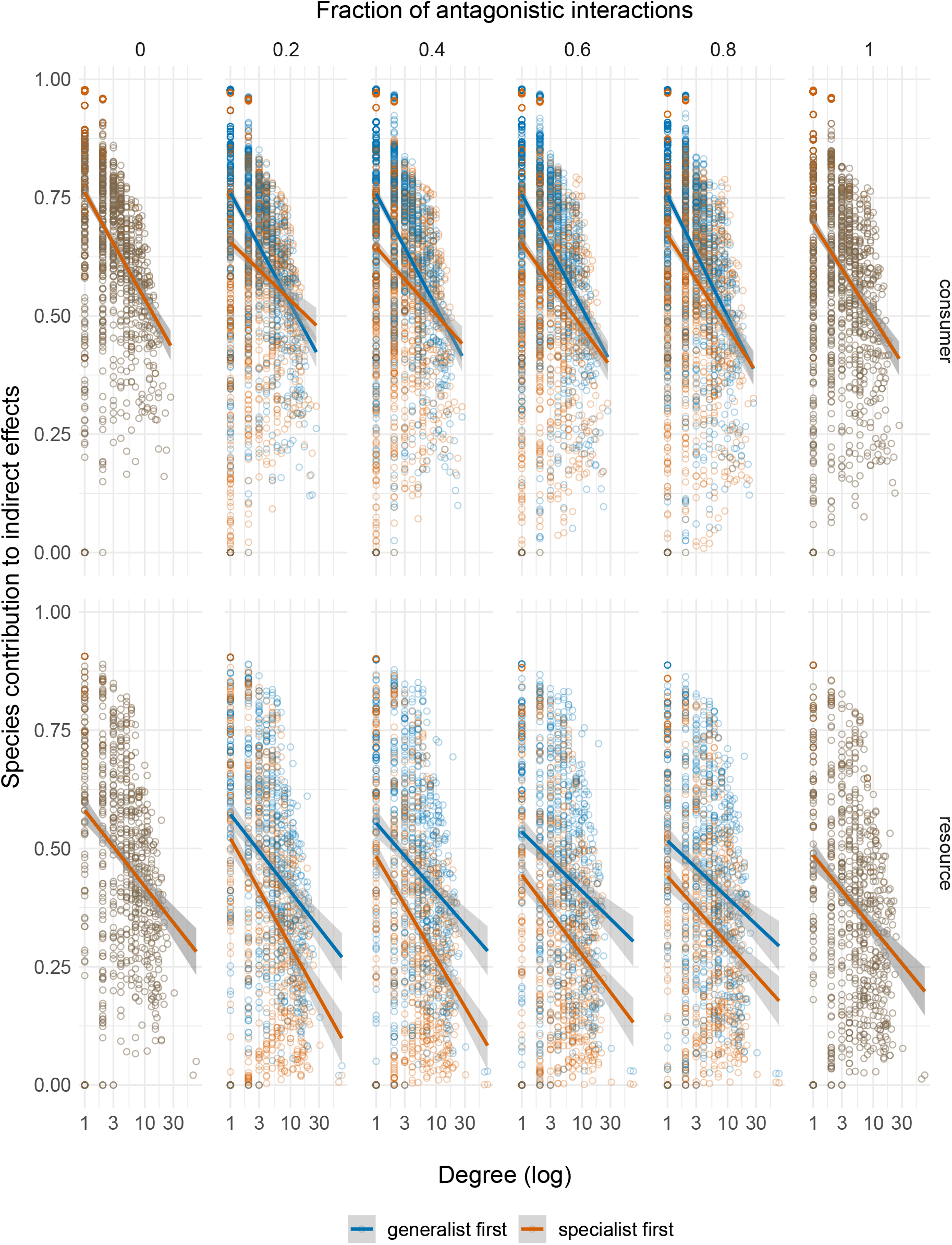
Effect of species degree on species contribution to indirect effects, along the transition from antagonism to mutualism. Numbers above the panels denote the fraction of antagonistic interactions in the networks. Colours represent the strategy used to convert antagonistic to mutualistic interactions. Lines highlight the linear relation between variables.

